# Use of Metagenomic Reference Materials to Compare Lysis and Extraction Methods Against a Known Input

**DOI:** 10.64898/2025.12.31.697245

**Authors:** Michael M. Weinstein, Braden Tierney, Elaine Wolfe, Shuiquan Tang, Venu Lagishetty, Jonathan P. Jacobs, Ryan Kemp, Christopher Mason

## Abstract

Inconsistent cellular lysis is a major source of inaccuracy in microbial analysis; there is a need to develop objective and standardized evaluation of lysis efficiency across diverse methods. Using a mock microbial community as a ground truth, we built a bioinformatic and experimental toolkit for this purpose, comparing the performance of chemical, thermal, enzymatic, and various physical lysis protocols. We evaluated over 150 lysis conditions and observed the highest performance from mechanically- and materially-optimized bead-based approaches. To extend these findings to a practical setting, we demonstrated that the Firmicutes to Bacteroides ratio, a frequently used compositional metric in gut microbiome analysis is biased heavily by insufficient lysis, indicating the importance of benchmarking ongoing methodological development.

## Body

The development of high-throughput, sequencing-based microbial community analysis has demonstrated the diversity and impact of microbiomes on human and planetary health. However, the multiplicity and inherent complexities of metagenomic processing workflows, particularly in obtaining unbiased lysis and extraction of nucleic acids (Angebault et al. 2018; Costea et al. 2017; Dopheide et al. 2019; Ezzy et al. 2019; Fiedorová et al. 2019; Han et al. 2019; Hart et al. 2015; Jiang et al. 2019; Kennedy et al. 2014; Mallott, Malhi, and Amato 2019; Martin-Laurent et al. 2001; Mattei et al. 2019; Pankoke et al. 2021; Pathirana et al. 2019; Sáenz et al. 2019; Vesty et al. 2017; Vishnivetskaya et al. 2014; Xue et al. 2018) has called into question the accuracy and comparability of studies using different methods. Inconsistencies in the performance of lysis protocols has created challenges in drawing conclusions between different research initiatives, such as the Human Microbiome Project (HMP) (Gevers et al. 2012) and Metagenomics of Human Intestinal Tract (MetaHIT) (Qin et al. 2010). While both studies agreed upon the predominant phyla in the gut microbiome, quantitative differences between the community compositions of their study populations could not be determined due to variation in extraction methods (Wesolowska-Andersen et al. 2014). One potential source of this and similar biases, is the challenge of biophysical challenges to lysis, where cells are recalcitrant to lysis due to their cell wall composition. (Ruggieri et al. 2016; Sohrabi et al. 2016; Yuan et al. 2012)

Historically, the alpha diversity (Menke et al. 2017) of complex reference samples, such as soil and feces (Angebault et al. 2018; Costea et al. 2017; Dopheide et al. 2019; Ezzy et al. 2019; Fiedorová et al. 2019; Han et al. 2019; Hart et al. 2015; Jiang et al. 2019; Kennedy et al. 2014; Mallott, Malhi, and Amato 2019; Martin-Laurent et al. 2001; Mattei et al. 2019; Pankoke et al. 2021; Pathirana et al. 2019; Sáenz et al. 2019; Vesty et al. 2017; Xue et al. 2018), have been used to evaluate lysis efficacy. However, the exact microbial composition of naturally-derived samples cannot be known and can vary sample-to-sample, making them suboptimal tools for evaluating lysis efficacy. It is imperative that methods be developed using (1) reference materials with precisely known inputs in conjunction with (2) bioinformatic packages that leverage prior knowledge of these materials 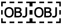 (Hornung, Zwittink, and Kuijper 2019; Lear et al. 2018; Ravel et al. 2014; Sinha et al. 2015; Stulberg et al. 2016; NIST 2019; Zengler et al. 2019)

Accordingly, as a tool to evaluate lysis efficacy on a known microbial community, we developed and present here the “Measurement Integrity Quotient” (MIQ) score and its accompanying bioinformatic workflows. In brief, the MIQ score divides the observed versus expected abundance of each member of a mock community and is converted to a percentage, indicating the degree to which expected relative abundance aligns with observed. Rather than calculating the residual relative to the theoretical 100 percent of expected, though; it is necessary to account for manufacturing tolerances in these reference materials. This results in calculating the residuals relative to a band around the theoretical ± tolerance. Finally, the mean of these residual percentages is subtracted from 100, meaning that a “perfect” MIQ Score of 100 indicates no deviations from expected composition that cannot be accounted for by manufacturing tolerance while decreasing scores indicate increasing deviation.

We then used the MIQ score to compare 161 common lysis and extraction kits and protocols available at the time this study was conducted. We observed a wide range of performances as indicated by MIQ Score (Fig. 1A). To isolate just the lysis method, beads from several different kits were used with an identical bead beating protocol and identical extraction methods, which yielded similar variations in performance, suggesting that the method of lysis was playing a major role in the variable performance of these methods (Fig. 1B).

**Figure 1.**
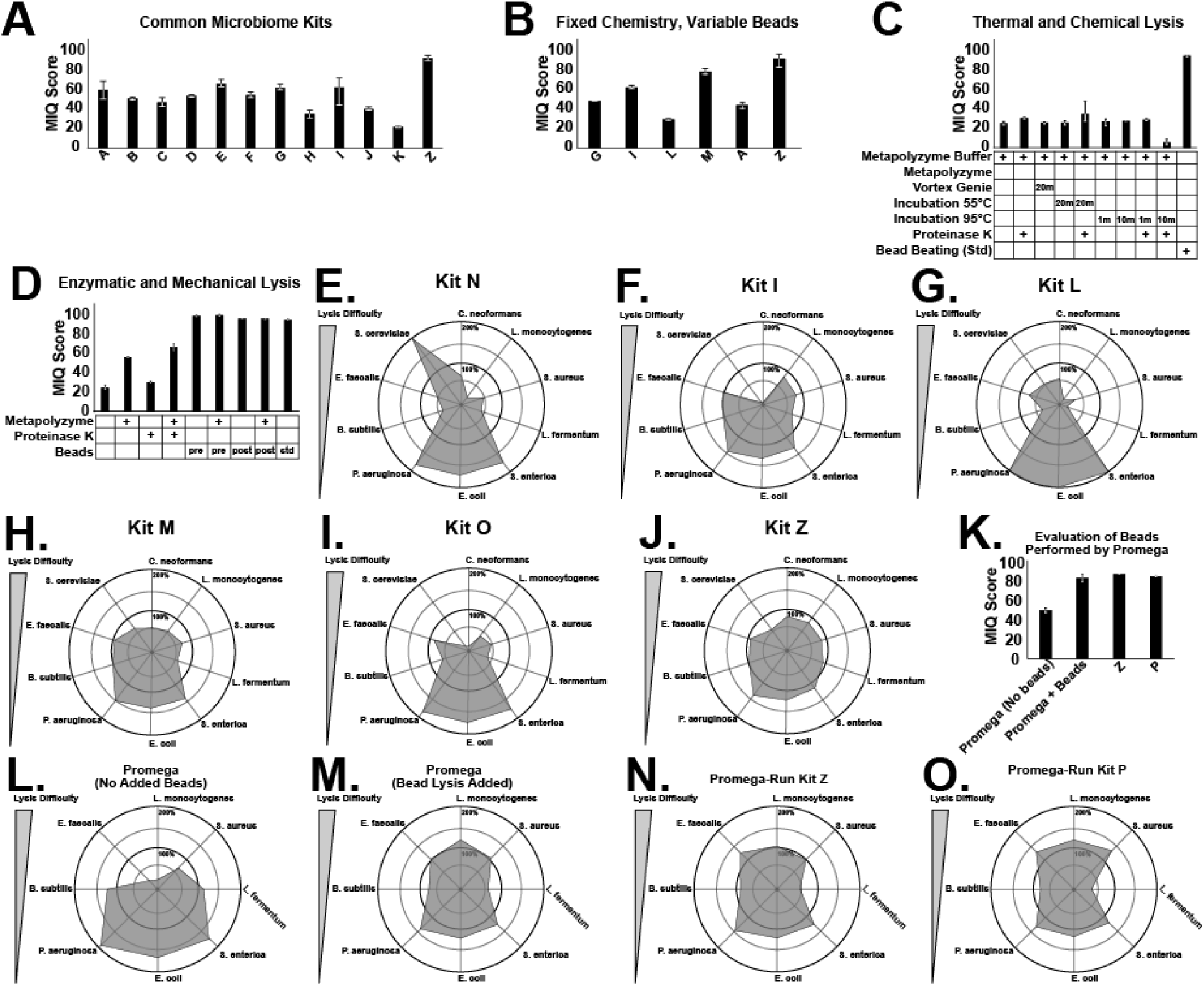
**A.** Comparing several commercially available microbiome kits available at the time of the study using their suggested lysis protocols with a minimum of 3 replicates per kit. A wide variation in performances as judged by the MIQ score could be observed (Error bars represent the observed range). **B.** This variability could be observed even when the lysis beads from multiple kits were used with the same post-lysis extraction and purification method, suggesting that the lysis mechanism contributed significantly to variation in performance. **C.** Applying this technique to various thermal protocols using ZymoBiomics’ protocol as a control demonstrated that no tested thermal lysis method gave high performance. **D.** Using the same method to test enzymatic and mechanical (bead-based) lysis showed improved performance over thermal lysis by enzymatic methods, but that a bead-based lysis method was both necessary and sufficient to give the maximum performance of all methods tested. **E-J.** Radar plots generated using the MIQ score bioinformatics package were capable of showing specific patterns of lysis deficiencies such as (**E**) irregular, inconsistent lysis, (**F**) near-total loss of fungal species, and (**G-J**) varying degrees of shift towards gram-negative microbes (all present at the bottom of the plot). **K**. Standard Promega lysis without beads showed a markedly diminished performance compared to protocols utilizing beads, including the standard Promega protocol with an added bead step. **L-O**. MIQ package radar plots showed a typical lysis bias pattern in data generated by Promega when beads were absent (L), while methods using beads showed improved accuracy and decreased lysis bias (M-O).

We next compared several thermal lysis methods (Fig. 1C) as well as enzymatic and mechanical (Fig. 1D) methods. None of the thermal lysis protocols we tested were able produce even moderately accurate results, but we did see significant improvement using enzymatic lysis methods. Bead-based lysis proved both necessary and sufficient for top performance among methods tested here. Radar plots (Fig 1E-J) revealed clear patterns on low-performing lysis methods. These approaches showed strong bias toward overrepresentation of gram-negative microbes, which is to be expected from incomplete lysis. This is concordant with historical observations of gram-negative enrichment in microbiome samples that were insufficiently lysed (Wesolowska-Andersen et al. 2014; Sohrabi et al. 2016; Yuan et al. 2012).

An independent team at a separate facility repeated this analysis and reproduced the findings. Specifically, they showed a similar pattern characteristic of poor lysis in the absence of beads (1K-L) that resolved with their introduction (1M). However, after lysis bias was addressed, we noted that *Lactobacillus* was still not detected at its expected abundance (1M-O).

To further optimize bead-based lysis, we tested several combinations of bead sizes and materials, while keeping all other portions of the analysis identical (Fig. 2A). We observed that zirconia-based beads of a size between 0.1 and 0.5mm gave superior performance to other bead compositions. However, the best overall lysis was accomplished by a mixture of 0.1 and 0.5mm bead materials.

**Figure 2.**
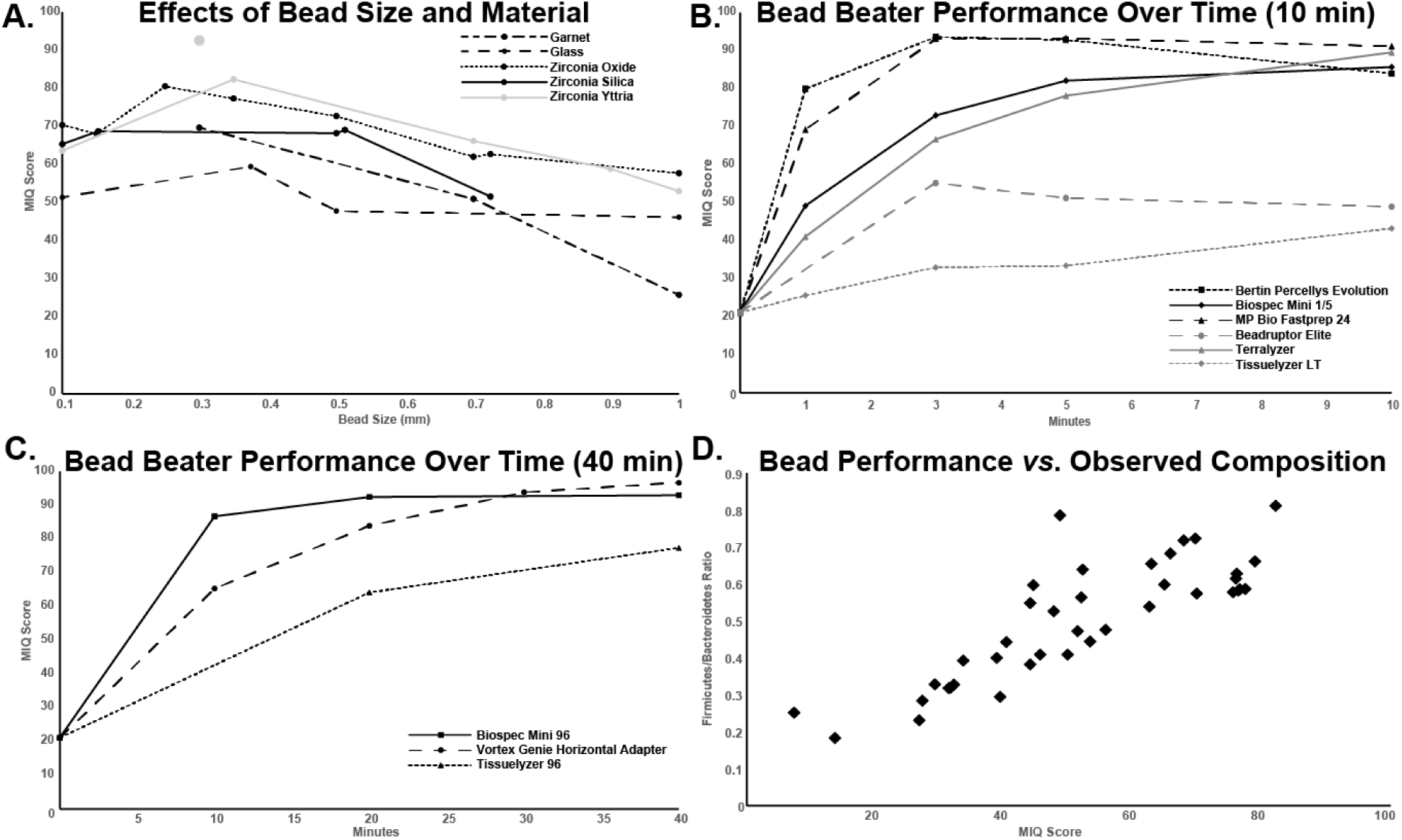
**A.** Testing varying combinations of bead material (line-color) and size (x-axis), optimal size and material combinations could be determined using the MIQ score where zirconium ceramic beads overall showed superior performance with beads in the 0.1-0.5mm size range giving optimal performance for any material. Combining 0.1 and 0.5mm zirconium beads provided superior performance compared to any material/size combination tested. **B-C.** Testing these mixed 0.1 and 0.5mm zirconium beads on different bead beaters for varying amounts of time showed clear patterns of increasing efficacy *vs*. time until a plateau was reached, indicating the maximum performance and optimal beating time for that specific unit. Units with larger motors were tested to 10 minutes (**B**) while smaller, less energetic units were tested out to 40 minutes (**C**). **D.** Comparing bacteria-only MIQ scores (x-axis) and observed Firmicutes/Bacteroidetes ratio on a repeated fecal sample (y-axis) on several different bead/material combinations (each point representing the mean of at minimum 3 replicates) reveals a strong correlation (Pearson *r*=0.835, *p*=3.7e-11) between the two values exhibiting the expected behavior of lower-performing beads increasing the apparent abundance of Firmicutes while beads with increasing performance characteristics against a known input standard exhibit an increased apparent abundance of tough-to-lyse Firmicutes.

We additionally optimized (i.e., found conditions that maximized the MIQ score) bead beating protocols along the axes of runtime and specific bead beater (Figs. 2B and 2C). We found that no one bead beater was “the best,” but rather that optimal runtime varied for different devices. For example, MPBio FastPrep required 3 to 5 minutes to achieve MIQ scores of >90, whereas the Vortex Genie required 30 to 40 minutes for the same result.

We additionally validated these findings regarding bead beating protocol optimization with another independent collaborator (the UCLA Microbiome Core). They compared their internal lysis pipelines, which had variable performance from sample to sample, to the protocols developed in this study (Fig. S1). Using a Vortex Genie-based lysis protocol, they observed a rapid increase in accuracy, as judged by MIQ score calculation.

Finally, to assess the potential “real-world” consequences of lysis biases, we compared MIQ scores on several bead material/size combinations with the Firmicutes/Bacteroidetes ratio (F/B ratio) generated on a repeated, preserved fecal sample (Fig. 2D). There was a strong positive correlation between increasing MIQ score and increasing observation of firmicutes, suggesting that technical variations can have a profound effect on the observed microbial community composition. In other words, the F/B ratio cannot be considered a reproducible metric for microbiome analysis without considering lysis strategy. While this observation may cause concern for the integrity of historical results, it is important to remember that differences observed using a biased method can still be valid, so long as the bias is consistent in behavior from sample to sample and run to run.

## Methods

### Measurement Integrity Quotient (MIQ) Score calculation

MIQ score source code can be found online for both 16S (https://github.com/Zymo-Research/miqScore16SPublic) and whole genomic (shotgun) sequence data (https://github.com/Zymo-Research/miqScoreShotgunPublic).

### Comparison of Commercial Kits and Laboratory Protocols

Commercial kits and popular DNA extraction methods at the time of analysis were tested on ZymoBIOMICS Microbial Community Standard (Zymo Research, Catalog # D6300) and stool samples, designated stool sample #98, in triplicate exactly according to manufacturer-supplied protocols. Notable differences in lysis conditions of each protocol that was performed according to manufacturer specifications are listed in the table below.

| Protocol | Beads | Bead Beating | Chemical/Enzymatic | Chemical/Thermal |
| --- | --- | --- | --- | --- |
| ZymoBIOMICS DNA MiniPrep | 0.1mm/0.5mm zirconia (0.6mL dry volume) | Bertin Precellys, 9000rpm, 4 x 60 sec | None | None |
| IHMS Protocol Q | 0.1mm zirconia (0.3g) | MP FastPrep-24, 6.5m/s, 8min 15 sec (1 minute on, 5 minutes off) | PK + AL buffer (70C, 10min)* | ASL buffer (95C, 15min)* |
| HMP Protocol | garnet beads | Vortex Genie, max speed, 10 minutes | None | PowerBead solution 65C for 10 min, followed by 95C for 10 min |
| DNeasy PowerSoil | garnet beads | Vortex Genie, max speed, 10 minutes | None | None |

\*For the IHMS, ASL buffer incubation is prior to bead beating, PK + AL buffer is after bead beating.

### Comparison of Commercial Kits and Laboratory Protocols with Standardized Bead Beating Parameters

Commercial kits and popular DNA extraction methods were tested on ZymoBIOMICS Microbial Community Standard (Zymo Research, Catalog # D6300) and stool samples, designated stool sample #98, in triplicate exactly according to manufacturer-supplied protocols using a standardized bead beating procedure. For all kits where bead beating was used as the lysis method, bead beating was carried out using the Scientific Industries Vortex Genie with Horizontal-(24) Microtube Adaptor with a maximum of 18 samples loaded per run unless otherwise specified.

| **Manufacturer** | **Kit or Protocol** | **Catalog #** |
| --- | --- | --- |
| Qiagen | DNeasy PowerLyzer | 12855-100 |
| Qiagen | DNeasy PowerSoil* | 12888-100 |
| N/A | HMP Protocol | N/A |
| N/A | IHMS Protocol Q | N/A |
| Sigma-Aldrich | Metapolyzyme protocol | MAC4L |
| Machery-Nagel | Nucleospin Soil | 740780.50 |
| Omega Bio-Tek | E.Z.N.A.® Soil | D5625-01 |
| Invitrogen | Purelink™ Microbiome | A29790 |
| Qiagen | QIAamp® PowerFecal | 12830-50 |
| Qiagen | QIAamp® DNA Stool | *Discontinued* |
| Zymo Research | ZymoBIOMICS DNA Miniprep | D4300 |

\*Formerly distributed as MOBIO's PowerSoil kit.

### Mechanical Lysis: Comparisons of Lysis Matrices

Identical conditions were used for all samples. Each lysis matrix used for mechanical lysis was tested in triplicate at a minimum. For each condition, 0.6 ml of beads were dispensed into 2 mL tubes. For conditions where lysis matrices were mixed, a volume/volume ratio was used. After sample addition to the lysis tube, the total volume was raised to 950 uL using ZymoBIOMICS Lysis Solution (Zymo Research, Catalog # D4300-1-150). Mechanical agitation was performed using the Bertin Precellys Evolution at 9000 rpm, cycling between 60 seconds of mechanical lysis and 120 seconds of rest for 4 cycles, for a total of 4 minutes of mechanical lysis.

After mechanical lysis, the ZymoBIOMICS DNA Miniprep Kit (Zymo Research, Catalog # D4300) was used for the DNA purification of all samples according to the manufacturer’s protocol, beginning immediately after the lysis step. When lysis matrices from commercially available kits were compared, the above DNA purification was also used in order to isolate the mechanical lysis as a variable. Where a repeated fecal sample was used, the sample was preserved frozen at 10% W/V in a solution of 15% glycerol in phosphate buffered saline (PBS). Note, the lysis matrix from the ZymoBIOMICS DNA Miniprep Kit was used as an internal control to ensure replicability between extractions, resulting in multiple sets of triplicates.

### Mechanical Lysis: Comparison of Bead Beaters

Samples were placed into a ZymoBIOMICS BashingBead Lysis Tube containing a mix of 0.1 and 0.5mm yttria zirconia beads with the final volume being raised to 950 uL using ZymoBIOMICS Lysis Solution (Zymo Research, Catalog # D4300-1-150). Mechanical lysis was performed using different mechanical lysis devices over a titration of time. Each condition was performed in triplicate at a minimum. The same lysis matrix was used for each mechanical lysis device condition in order to isolate the effect of the mechanical lysis device.

The more energetic bead-beaters (such as Precellys and FastPrep) were cycled on and off to allow for the dissipation of built-up heat generated by the bead collisions in the sample while less energetic systems (such as the vortex genie) lowered heat generation during lysis enough that heat dissipation from the tube surface during lysis was adequate to prevent sample breakdown and tube rupture.

After mechanical lysis, the ZymoBIOMICS DNA Miniprep Kit (Zymo Research, Catalog # D4300) was used for the DNA purification of all samples according to the manufacturer’s protocol, beginning at step 3. Following extraction, library preparation, and sequencing using the methods described in this section, accuracy of the extraction was measured using the MIQ system.

### Chemical/Enzymatic Lysis

Metapolyzyme (Sigma-Adrich, Catalog # MAC4L) in combination with a sodium dodecyl sulphate rich buffer, Solid Tissue Buffer (Zymo Research, Catalog # D4086-2-22), proteinase K, and/or bead beating were used for lysis of ZymoBIOMICS Microbial Community Standard (Zymo Research, Catalog # D6300) and stool sample #93 in triplicate. The following condition combinations for lysis were tested:

Case 1: Metapolyzyme
Case 2: Metapolyzyme followed by Solid Tissue Buffer + proteinase K
Case 3: Bead beating followed metapolyzyme
Case 4: Metapolyzyme followed by bead beating
Case 5: Bead beating only control

Metapolyzyme was used according to manufacturer instructions were incubated with Metapolyzyme for 4 hours at 35C. For case 2, Solid Tissue Buffer and proteinase K were added afterward, then incubated for 1 hour at 55C. For cases 3-5, bead beating was performed using the MP FastPrep-24 at 6.5 m/s, 1 minute on and 5 minutes off, for 5 cycles. For every experimental condition except for case 5, sample sets were also run with PBS blanks in place of Metapolyzyme as negative controls. After lysis, samples were purified using ZymoBIOMICS DNA Miniprep Kit (Zymo Research, Catalog # D4300), starting at step 3 of the manufacturer supplied protocol.

### Chemical/Thermal Lysis

Triplicate samples of ZymoBIOMICS Microbial Community Standard (Zymo Research, Catalog # D6300) and fecal sample #98 were lysed using a combination of Solid Tissue Buffer and proteinase K incubated at varying temperatures and time, and/or bead beating.

Case 1: Solid Tissue Buffer (55C, 20 min)
Case 2: Solid Tissue Buffer + proteinase K (55C, 20 min)
Case 3: Solid Tissue Buffer and bead beating (25C, 20 min)
Case 4: Solid Tissue Buffer (95C, 1 min)
Case 5: Solid Tissue Buffer (95C, 1 min) + proteinase K (55C, 20 min)
Case 6: Solid Tissue Buffer (95C, 10 min)
Case 7: Solid Tissue Buffer (95C, 10 min) + proteinase K (55C, 20 min)

Bead beating for case 3 was performed using the Vortex Genie at maximum speed at room temperature. All samples were purified after lysis using ZymoBIOMICS DNA Miniprep Kit (Zymo Research, Catalog # D4300) starting at step 3 of the manufacturer-supplied protocol.

### Additional Comparison of Commercial Kits and Extraction Protocols

Multiple commercially available kits and widely used DNA extraction protocols were compared using the ZymoBIOMICS Gut Microbiome Standard (Zymo Research, Catalog # D6331) to ensure reproducibility of the results observed with the ZymoBIOMICS Microbial Community Standard (Zymo Research, Catalog #D6300). The HMP Protocol, IHMS Protocol, DNeasy PowerSoil Kit (Qiagen, Catalog # 12888-100), DNeasy PowerSoil Pro Kit (Qiagen, Catalog # 47016), and ZymoBIOMICS DNA Miniprep Kit (Zymo Research, Catalog # D4300) were used according to manufacturer-supplied protocols to extract triplicate samples.

Detailed protocols used in this study, including ones from collaborating labs, can be found in Appendix 1.

### Library Preparation

After DNA Purification, DNA was quantified using UV absorbance via Nanodrop. Samples with A260/280 or A260/230 ratios below 1.8 were cleaned using the Select-a-Size DNA Clean & Concentrator (Zymo Research, Cat. # D4084). Prior to library preparation, DNA was also quantified via Qubit dsDNA HS Assay Kit (Thermo Fisher), or the Qubit dsDNA BR Assay Kit (Thermo Fisher) in instances when the sample was out of range of the HS kit.

Sequencing libraries were prepared from microbial DNA using the KAPA HyperPlus Library Preparation Kit (Kapa Biosystems, Wilmington, MA) with up to 100 ng DNA input following the manufacturer’s protocol using internal single-index 8 bp barcoding sequences and TruSeq Adapters (Illumina, San Diego, CA).

### Library Preparation at 3^rd^ Party Testing Site (Promega)

The DNA purified from the ZymoBIOMICS Microbial Community Standard was quantified using QuantiFluor® ONE fluorescent dye, and samples were diluted to a final concentration of 1ng/µl prior to sequencing. The sequencing protocol used was an adaptation of the 16S Metagenomic Sequencing Library Preparation (“MiSeq System Denature and Dilute Libraries Guide,” n.d.). The amplicon reaction was cleaned up using ProNex® Size-Selective Purification System with a 1.2:1 ratio of ProNex® Chemistry to sample volume and eluted in a volume of 40µl. Indexing reactions were performed with GoTaq® Long PCR Master Mix and Nextera XT dual-indexing primers. Reactions were again cleaned up with a 1.2:1 ratio of the ProNex® Size-Selective Purification System chemistry and assay and pooled at equimolar volumes before denaturing and diluting according to the Illumina MiSeq System Denature and Dilute Libraries Guide. The library pool was combined with 5% PhiX prior to sequencing on the MiSeq system at 2 x 300bp with the MiSeq Reagent Kit v3. The data were analyzed using a Mothur (Schloss et al. 2009) based bioinformatics pipeline.

---

**Figure S1:**
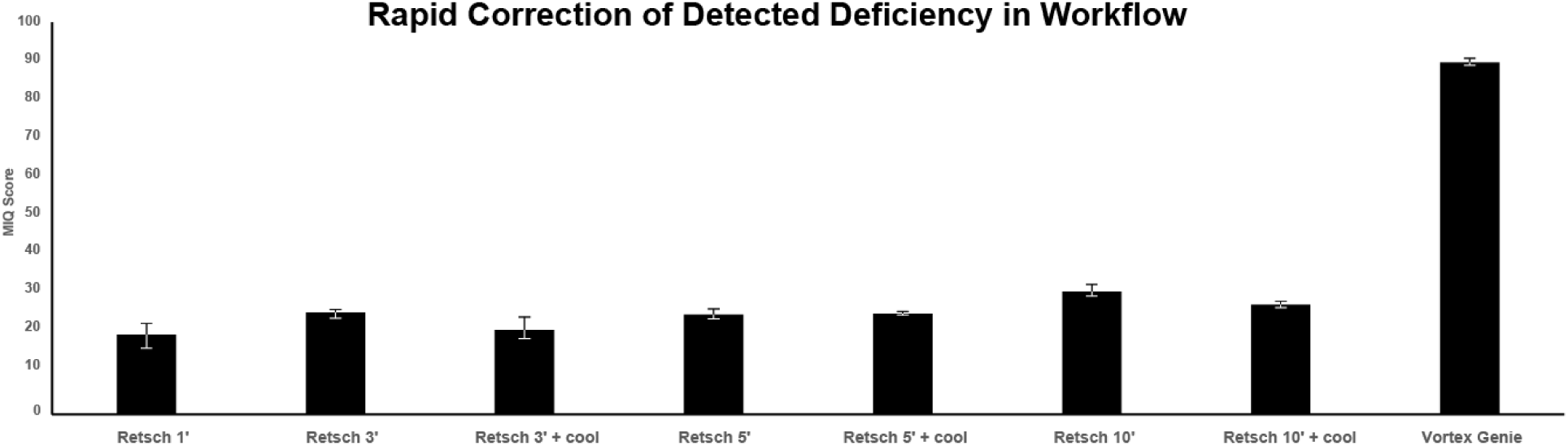
The UCLA Microbiome Core, upon benchmarking their primary lysis method (utilizing a Retsch bead beater) noticed low overall accuracy from their workflow despite several different lysis protocol variations. Upon swapping out the Retsch-based method for a Vortex Genie-based method that had previously demonstrated consistently high performance, the deficiencies in their accuracy scores rapidly resolved.

## Notes

### Competing Interest Statement

Zymo Research and Promega both manufacture microbiome sample preparation kits and reagents. Authors affiliated with these entities have this status indicated in their information.

